# Mitochondrial dysfunction: unraveling the elusive biology behind anti-TNF response during ulcerative colitis

**DOI:** 10.1101/2024.06.18.599496

**Authors:** Dimitrios Kioroglou, Ainize Peña-Cearra, Ana M. Corraliza, Iratxe Seoane, Janire Castelo, Julian Panés, Laura Gómez-Irwin, Iago Rodríguez-Lago, Jone Ortiz de Zarate, Miguel Fuertes, Itziar Martín-Ruiz, Monika Gonzalez, Ana Mª Aransay, Azucena Salas, Héctor Rodríguez, Juan Anguita, Leticia Abecia, Urko M. Marigorta

## Abstract

**Background:** Recent studies hint at mitochondrial genes influencing UC patient response to anti-TNF treatment. We evaluated this hypothesis by following a targeted strategy to identify gene expression that captures the relationship between mitochondrial dysregulation and response to treatment. Our objective was to initially examine this relationship in colon samples and subsequently assess whether the resulting signal persists in the bloodstream.

**Methods:** We analyzed the transcriptome of colon samples from an anti-TNF treated murine model characterized by impaired mitochondrial activity and treatment resistance. We then transferred the findings that linked mitochondrial dysfunction and compromised treatment response to an anti-TNF treated UC human cohort. We next matched differential expression in the blood using monocytes from peripheral blood of controls and IBD patients, and we evaluated a classification process at baseline with whole blood samples from UC patients.

**Results:** In human colon samples, the derived gene-set from the murine model showed differential expression, primarily enriched metabolic pathways, and exhibited similar classification capacity as genes enriching inflammatory pathways. Moreover, the evaluation of the classification signal using blood samples from UC patients at baseline highlighted the involvement of mitochondrial homeostasis in treatment response.

**Conclusion:** Our results highlight the involvement of metabolic pathways and mitochondrial homeostasis in determining treatment response and their ability to provide promising classification signals with detection levels in both colon and bloodstream.

## Introduction

Ulcerative colitis (UC) is an inflammatory bowel disease (IBD) of increasing worldwide incidence, characterized by relapsing and remitting periods. Lacking a definitive cure, the focus of treatments is to alleviate and control disease symptoms. Although current anti-TNF treatments can extend the remission periods, a considerable 10-30% of patients do not respond to the initial treatment, or lose their ability to respond (23-46%) over time [1].

Through comparisons of gene expression profiles of responder and non-responder patients, several genes with biomarker potential have been proposed. This includes intestinal Oncostatin M (OSM), IL13RA2 and TREM1, which were first identified using mucosal samples [2–4]. Similarly, Haberman *et al.* [5] provided a gene signature derived from rectal biopsies at baseline. Although originally described for response to corticosteroids, the gene signature had a similar efficiency on a cohort that received anti-TNF treatment, whereas reduced mitochondrial biogenesis was reported as an aggravating factor.

To avoid the invasive nature of the patient’s monitoring via biopsies, identifying whole blood biomarkers is a priority. For instance, TREM1 expression is arising as a promising predictive biomarker, although the evidence is not yet conclusive [4,6]. In addition, a qPCR classifier aimed at classifying patients according to disease course is available [7].

In spite of the valuable endeavors towards classification solutions, current methodologies are based on differential gene expression analysis that lacks a perturbation source. To derive insights on the underlying mechanism and provide a gene signature that includes a reasonable number of differentially expressed genes (DEGs), studies resort to pathway enrichment analysis. However, this approach is based on using databases that may present limitations, including outdated or missing pathway information, and, most importantly, they are not disease specific [8–10]. Thus, the accuracy of the emerging patterns is uncertain and may result in lack of clinical utility for guiding therapy, as has recently been shown by Noor et al. in a study of Crohn’s disease where the authors evaluated the classification efficiency of a blood-based biomarker [11]. Thus, more refined and targeted methodologies are necessary for advancing biomarker discovery.

Drug response in the context of UC involves widespread changes in inflammation, and as a result pathways with significant contribution to the treatment response can be overshadowed by inflammatory pathways. In order to unravel the significant contribution of obscured biology, our approach utilizes a murine model coupled with a perturbation source to identify explicit gene expression patterns, and capture underlying mechanisms that represent promising candidates to decipher the lack of response to treatment.

Recent evidence has highlighted the potential involvement of mitochondrial dysfunction in disease severity and treatment response [5], emphasizing the need to identify underlying factors that contribute to mitochondrial dysregulation. In this regard, we have previously described that the absence of Methylation-controlled J protein (MCJ), or DnaJC15, an endogenous negative regulator of mitochondrial complex I, leads to metabolic alterations, increases respiration and adenosine triphosphate (ATP) synthesis and stimulates the formation of respiratory supercomplexes, limiting oxidative stress [12–14]. This qualifies MCJ as a good perturbation source to simulate complex I mitochondrial dysfunction. Our goal was to transfer the findings to human cohorts and assess the persistence of the emerged patterns.

In the context of experimental colitis in a murine model, we have previously shown that deficiency of MCJ contributes to disease severity by modulating the gut microbial composition of the host [15–17]. Moreover, recently we have reported the critical role played by macrophage mitochondrial function in the gut ecological niche, which can substantially affect the severity of inflammation and the ability to successfully respond to anti-TNF therapies [18]. The lack of response to the treatment was expressed as an aggravated phenotypic profile, where treated MCJ-deficient mice displayed significantly increased weight loss and histological damage compared to the treated wild-type mice.

In the current study, we profiled the transcriptome of colonic tissue from the aforementioned MCJ-deficient murine model of DSS-induced colitis [18], to generate a gene set that captures the relationship between mitochondria dysfunction and lack of response to anti-TNF treatment. Activated monocytes are recruited to the site of inflammation induced by DSS where they differentiate into macrophages. The latter can be polarized towards different subpopulations that can initiate either pro-inflammatory or anti-inflammatory responses [21]. It has been postulated that the FcγR signaling pathway is implicated in the modulation of these diametrically opposed responses [22]. Moreover, it has been suggested that the formation of immune complexes between soluble TNF and therapeutic anti-TNF compounds is necessary for the FcγR-mediated suppression of pro-inflammatory cytokines. We have shown that MCJ expression is restricted to macrophages in colonic tissue, and MCJ-deficiency leads to upregulation of genes belonging to the FcγR signaling pathway and aggravation of the DSS-induced pathology [18]. Thus, we hypothesize that mitochondrial dysregulation can indirectly compromise macrophage function and especially their ability to initiate inflammation-resolving responses.

As summarized in Figure 1, we followed a progressive strategy where the captured gene set from the murine model was initially transferred and evaluated in a human cohort that provided transcriptomic data of colonic tissue. Subsequently, we explored whether the captured signal can be detected in blood samples by applying a successive refinement with two human cohorts profiling the transcriptome of peripheral blood monocytes, and whole blood samples. Finally, we utilized an external dataset of whole blood samples from UC patients to evaluate the replication of the findings in blood.

**Figure 1:**
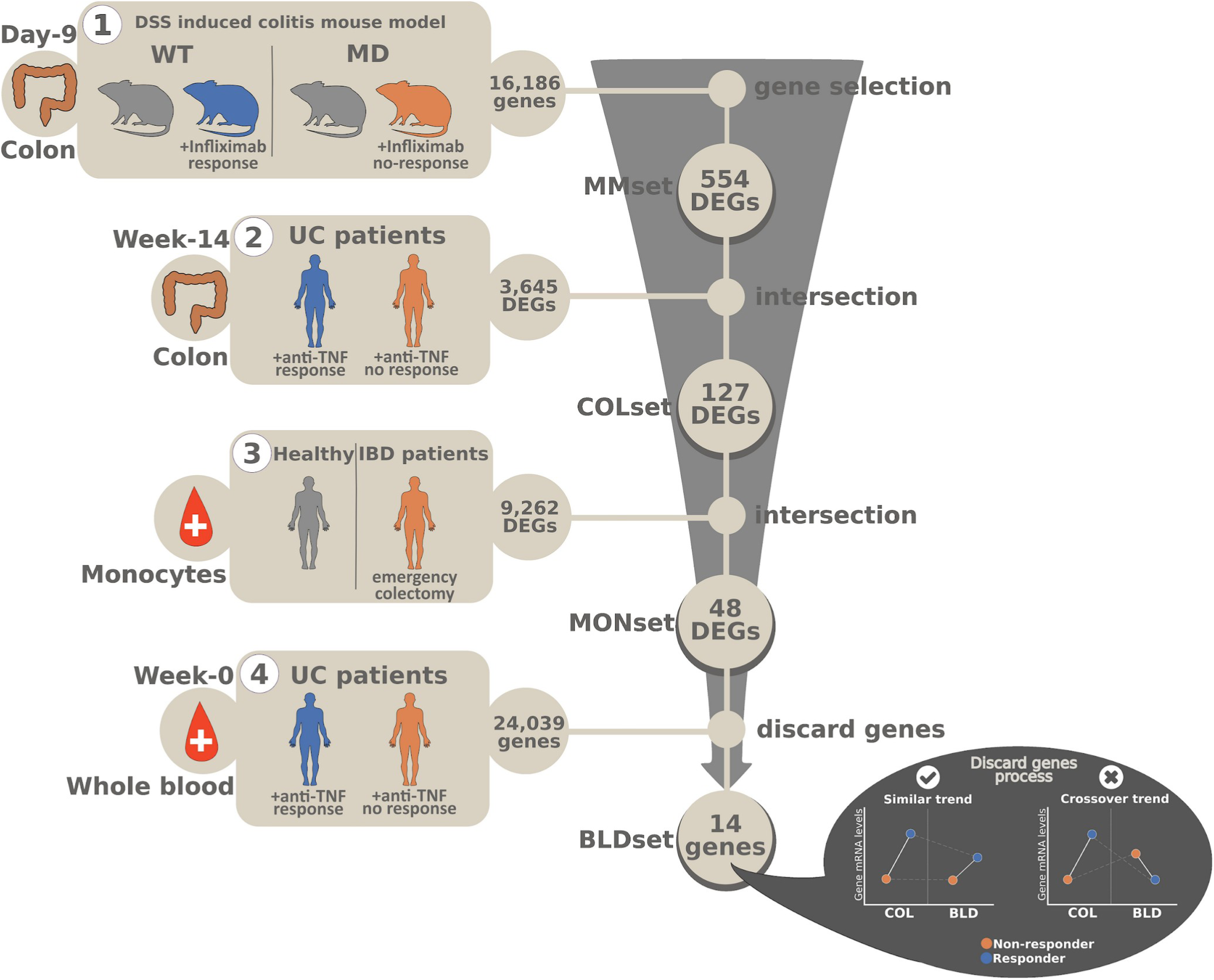
Experimental design. **Phase 1**: The analysis was based initially on a murine model of experimental DSS induced colitis characterized by MCJ-deficiency (MD) and mitochondrial complex I dysfunction. A set of 554 genes (MMset) was selected that showed strong relation with mitochondrial genes and response to anti-TNF treatment. **Phase 2**: We transferred the captured signal to a human cohort of UC patients that was treated with anti-TNF and provided colon samples. We identified a set of 127 genes (COLset) that match differential expression in MMset. **Phase 3**: We identified a classification signal of 48 genes (MONset) from the COLset that can be detected in circulating monocytes by comparing healthy individuals and IBD patients with an aggravated state of the disease that proceeded to emergency colectomy. **Phase 4**: Using whole blood samples of UC patients before anti-TNF treatment, we identified a signal of 14 genes from MONset (BLDset). The 14-gene based BLDset became the input of a classification process regarding response before anti-TNF treatment.

This is the first study to use a perturbation source that simulates mitochondrial dysregulation to scrutinize the involvement of the latter on the lack of response to anti-TNF treatment during ulcerative colitis in human patients. Our findings shed new light on the role of metabolic pathways and mitochondrial homeostasis in determining treatment response, and pave the way for further research into the mechanisms underlying drug response in ulcerative colitis and other inflammatory diseases.

## Methods

### Ethics

All procedures involving animals, including their housing and care, were carried according to the guidelines of the European Union Council (Directive 2010/63/EU) and Spanish Government regulations (RD 53/2013), and with the approval of the ethics committee of CIC bioGUNE (Spain; permit number CBBA-0615) and the Competent Authority (Diputación de Bizkaia). All human samples were processed within the context of the IBD Sample Collection registered at the HCB-IDIBAPS Biobank and the protocol was approved by the Institutional Ethics Committee of the Hospital Clinic de Barcelona (HCB/2023/0523). Additional approval for the human samples was provided by the Clinical Research Ethics Committee from Euskadi (CEIC-E) following the Helsinki convention (PI2016034). In all human cohorts, patients provided written informed consent following extensive counseling.

### Animal model and experimental design

The murine model [18] included MCJ-deficient (MD) and wild-type (WT) male mice and is referred to as MM dataset herein. To evaluate the response to anti-TNF treatment, colitis was induced in WT and MD mice by administering dextran sodium sulfate (DSS) [15]. The whole experimental period lasted nine days and the mice were divided into two groups per genotype. One group received Infliximab (IFX^+^) treatment on the third and sixth day (WT-IFX^+^, MD-IFX^+^), whereas the second group did not receive the treatment (WT-IFX^-^, MD-IFX^-^). The distribution of mice per condition was 5 WT (2 WT-IFX^-^ and 3 WT-IFX^+^) and 5 MD mice (2 MD-IFX^-^ and 3 MD-IFX^+^). Colon samples were taken from all mice at the end of the experimental period, namely nine days after the treated mice had been administered Infliximab (day-3 and 6), and used to study RNA expression.

All procedures and specifications involving animal treatment and the design of the mouse model are described in the supplementary methods (S1).

### Human cohorts

Human cohorts are referred to with the abbreviations COL, MON and BLD. The COL cohort (N=28) included colonic biopsies at week-14 after anti-TNF treatment of UC patients. The MON cohort (N=31) included healthy blood donors and IBD patients that provided monocytes from peripheral blood. The IBD patients were individuals that presented an aggravated state of the disease and either did not respond to previous treatments or manifested severe symptoms. As a result, the IBD patients proceeded to emergency colectomy where blood samples were taken prior to operation. The BLD cohort (N=34) included whole blood samples from UC patients prior to anti-TNF treatment (week-0) and their response was evaluated at week-14. The COL and BLD cohorts provided in total 39 patients, from which 23 patients provided whole blood samples at week-0 and colon samples at week-14, whereas 5 patients provided only colon samples at week-14 and 11 patients only whole blood samples at week-0. Moreover, response to treatment was based on endoscopic activity. Age and gender of patients at inclusion are shown in Table 1 for each cohort. Cohort details and specifications regarding sample collection and treatment are provided in the supplementary methods (S2).

**Table 1:** Human cohorts specifications. Age is reported based on average and observed minimum and maximum values. Information regarding average age and gender was not available for the healthy individuals in the MON cohort.

|  | N | Age (MIN-MAX) | Gender (F/M) |
| --- | --- | --- | --- |
| COL cohort |  |  |  |
| responders UC | 7 | 41 (20-60) | 3/4 |
| non-responders UC | 21 | 46 (27-70) | 9/12 |
| MON cohort |  |  |  |
| healthy | 16 | N/A (18-65) | N/A |
| IBD patients | 15 | 49 (23-77) | 4/11 |
| BLD cohort |  |  |  |
| responders UC | 9 | 49 (24-87) | 3/6 |
| non-responders UC | 25 | 53 (32-78) | 11/14 |

### RNA extraction, preparation and quantification

In the mouse model and human cohorts, gene quantification was performed with RNASeq using the GRCm38 mouse reference and the GRCh38.p13 human reference, respectively. A gene was characterized as differentially expressed (DEG) for a given condition contrast, if it met the following criteria: FDR-corrected p-value (p-adj) *≤* 0.05 and absolute fold-change |*FC*| *≥* 1.5. The derived DEGs were later utilized for enrichment analysis from which only statistically significant enriched pathways were considered (p-adj *≤* 0.05). From each human cohort, the set of significant genes that was selected for downstream analysis was named as COLset, MONset and BLDset denoting the cohort of origin.

Detailed specifications regarding RNA extraction and preparation protocols, as well as RNASeq preprocessing and enrichment analysis, are given in the supplementary methods (S3).

### Incorporation of the Haberman et al. corticosteroid gene signature into the analytical pipeline

We incorporated into the analysis the gene signature provided by Haberman et al. [5]. The signature contains 115 genes that resulted from baseline rectal biopsies of 206 pediatric UC patients whose remission status was evaluated on the week-4 of corticosteroid treatment. Notably, this signature exhibited similar classification efficiency in a cohort treated with anti-TNF treatment. From the provided 115 genes, we selected 89 genes that were differentially expressed in our COL cohort. We named the selected genes as HABset, and were used to conduct a comparative assessment with the COLset on their ability to differentiate between responders and non-responders in the COL cohort.

### Replication of the findings in the colon with external datasets

To replicate the signal captured by COLset, we incorporated the gene expression datasets provided by two studies of Arijs *et al.* with accession number GSE16879 [19] and GSE73661 [20]. Both studies included responders (N=8, N=8 respectively) and non-responders (N=16, N=15 respectively) to anti-TNF treatment UC patients, with intestinal mucosal biopsies being collected at baseline and after four to six weeks after the treatment. Gene expression was quantified with Affymetrix Human Genome U133 Plus 2.0 and Affymetrix Human Gene 1.0 ST arrays, and the signal intensities were GCOS-normalized and log2 RMA-normalized respectively. To identify DEGs, we fitted a linear model using the LIMMA package of R, which additionally applied an empirical Bayes smoothing to the standard errors, and FDR-corrected the resulting p-values.

### Evaluation of the persistence of the captured signal from colon tissue in blood samples

Since MCJ expression is restricted in macrophages in colonic tissue [18], we sought to explore whether the captured signal from the COL cohort can be detected in the blood. To do so, we initially identified 48 genes (MONset) in the MON cohort that can distinguish IBD patients from non-IBD individuals. Later, we utilized whole blood samples before treatment from the BLD cohort, where our process for resolving crossover trends between the BLD and MON cohorts provided 14 genes (BLDset) that can classify responders and non-responders IBD patients to treatment. The process followed for the construction of the BLDset is described in the supplementary methods along with the concept of trends (S4).

For the classification process in the BLD cohort, we initially regressed-out age and sex from the gene expression data. To avoid overfitting and multicollinearity, we performed PCA on the gene expression and based a non-regularized logistic regression model on the first two PCs with Scikit-learn [23]. One additional advantage of this strategy is that it enables the visual evaluation of the model’s performance. To mitigate the impact of the imbalanced conditions, the weights 0.3 and 0.7 were used for the non-responder and responder classes respectively. To evaluate the training process, we performed a 12-fold cross-validation with 3 repeats calculating at each fold the F1-score metric for the BLD training and BLD test-set. The F1-score was chosen since it is a well-suited metric for imbalanced categorical variables. It is interpreted as the harmonic mean of the precision and recall metrics and ranges between 0 and 1. Finally, estimation of the decision boundary standard error was based on a bootstrapped 95% confidence interval with 1000 iterations, sampling with replacement and sample size of 34.

### Replication of the findings in blood with an external dataset

To evaluate the replication of the findings in the whole blood, we downloaded the publicly available RNASeq data provided by Mishra *et al.* [24] (GSE191328). It is a blood-based dataset that underwent a similar preprocessing procedure as the BLD cohort, and included three independent cohorts of IBD patients: two that received Infliximab treatment (referred to as discovery and replication cohorts by the authors) and one Vedolizumab. The patients were followed longitudinally from baseline until week-14, where they were classified according to remission status. From the available cohorts, we considered only UC patients and focused on the following time-points: baseline, week-2, week-6 and week-14. The discovery cohort is referred to as IFX (N=9) and the replication cohort as REP (N=9), whereas the Vedolizumab cohort is referred to as VED (N=10) in the current study. To avoid possible integration batch effects and to evaluate patients heterogeneity, we kept the IFX and REP cohorts separate.

Prior to model validation, the genes *SLC18A1* and *PPARGC1A* were omitted from the BLDset since they were not quantified across all cohorts and timepoints. We also processed each cohort with DESeq2 stabilizing the gene expression variance with the dispersion function whose parameters were previously estimated with the BLD cohort without specifying patient response. Thus, no patient’s response information was introduced during the dispersion estimate that could bias the classification process. Then, each cohort was projected on the PCs that were extracted during the model’s training process and the sample scores from the resulting transformation were used as a test-set. Finally, the performance of the classification process in each cohort was assessed using the F1-score metric.

## Results

### 554 DEGs in the mouse model (MMset) capture the interaction between the absence of MCJ and the response to anti-TNF

First, we performed transcriptomic analysis of the colon tissue from a murine model of experimental colitis that included wild-type (WT) and MCJ-deficient (MD) mice. The mouse groups were further subdivided into treated with anti-TNF (WT-IFX^+^,MD-IFX^+^) and untreated mice (WT-IFX^-^,WT-IFX^-^). As mentioned above, response to anti-TNF treatment was previously evaluated [18]. Similarly to the WT-IFX^-^ and MD-IFX^-^ mice, the MD-IFX^+^ mice displayed an aggravated profile characterized by significantly higher weight loss and histological damage compared to WT-IFX^+^. Additionally, in the current study we observed high efficiency of the model, with an 8.4-fold and 11.3-fold changes in the mRNA levels of *Dnajc15* between the MD-IFX^-^ and WT-IFX^-^ and the MD-IFX^+^ and WT-IFX^+^ groups, respectively (Figure S1).

To evaluate group differences, we performed principal component analysis (PCA) using all 16,186 quantified genes. The MD-IFX^+^ mice had unique distinguished profiles compared with the other three groups, forming a separate cluster across PC1 (45% of the observed variance, Figure 2A, left plot). The differences between the other three groups appeared weaker, clustering mainly across PC2 (18.8% of the observed variance).

**Figure 2:**
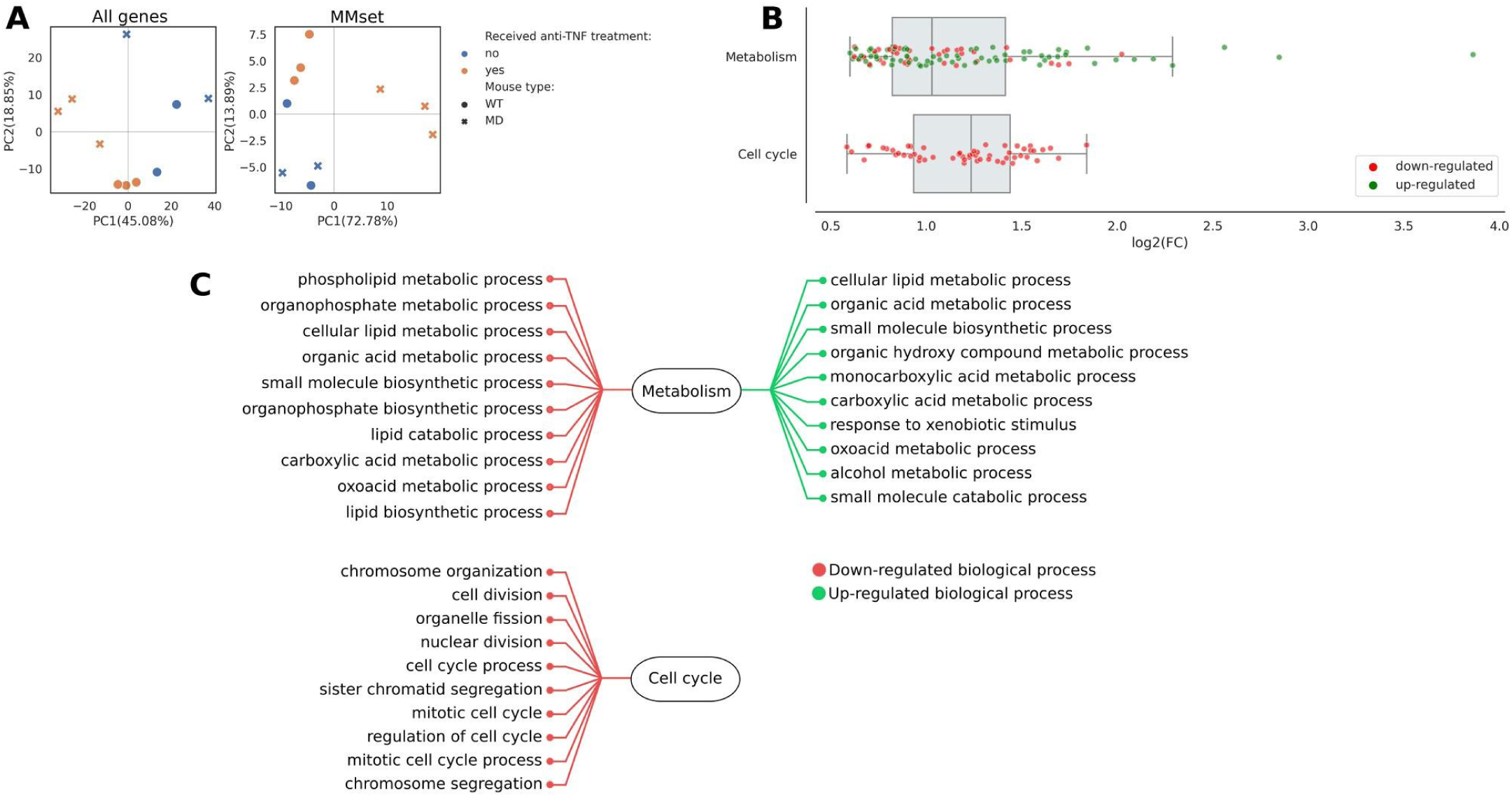
**A**) PCA using all 16,186 quantified genes (left) from the mouse model or only the MMset (right). **B)** Distribution of log2-fold changes (x-axis) between MD-IFX+ and WT-IFX+ for each enriched pathway based on MMset. In each pathway, up and down-regulated genes are indicated for the MD-IFX+ in relation to WT-IFX+ group. **C)** Up and down-regulated biological process related to each enriched pathway. Regulation is indicated for MD-IFX+ in relation to WT-IFX+ group.

Differential gene expression analysis corroborated these observations, with greater intra-group differences within the MD groups than among WT groups. The contrast between MD-IFX^+^ and MD-IFX^-^ resulted in 1,544 differentially expressed genes (DEGs), for only 12 DEGs in the WT-IFX^+^ and WT-IFX^-^ comparison. Of note, we observed an interaction between *Dnajc15* gene expression and the response to the anti-TNF treatment. Upon administration of Infliximab, 617 genes were differentially expressed between the treated groups (MD-IFX^+^ and WT-IFX^+^), for only 12 genes among the corresponding non-treated groups. These observations verify the utility of the mouse model, suggesting a significant interaction between *Dnajc15* expression and the response to Infliximab treatment.

We reasoned that the 617 DEGs from the MD-IFX^+^ versus WT-IFX^+^ comparison represent a set of genes with the capacity to decipher the mechanisms underlying the relationship between mitochondrial dysregulation and unresponsiveness to the treatment. After removing genes without corresponding human ENSEMBL identification, 554 DEGs were retained, referred to heretofore as the MMset (supplementary materials 1). As expected, PCA showed that the MMset recapitulates the differences between the MD-IFX^+^ and the other groups, forming a separate cluster across PC1 (72.78% of the observed variance, Figure 2A, right plot).

### MMset is associated with mitochondrial dysfunction

We used MitoMiner’s API [25] to identify genes located in the mitochondrion. We identified 50 genes from the MMset (supplementary materials 1). These genes had a median 1.21-fold average expression change between the non-treated (WT-IFX^-^ and MD-IFX^-^) groups, increasing to 1.86-fold change between the treated groups (Figure S2). These results reinforce the previous idea of an interaction between anti-TNF administration and mitochondrial dysfunction.

We identified other notable changes in genes with relevant roles in mitochondrial homeostasis. *Phgdh,* known to compromise the electron transport chain activity and lead to increased production of ROS [24], was significantly down-regulated in the MD-IFX^+^ group. We observed similar changes for *Hif1a* and *Ripk3*, that play a role in modulating mitochondrial function and metabolism [27,28]. In turn, we observed significant up-regulation of *Ppargc1a* and *Pink1*, which are involved in regulating mitophagy and mitochondrial biogenesis [29].

### The major enrichment in MMset concerned metabolic pathways

To further characterize the biological effects underlying the MMset, we performed enrichment analysis of the full set. Subsets of 107 and 56 genes were associated with metabolic and cell cycle pathways, respectively (supplementary materials 2). From the subset associated with metabolic pathways, 65.4% of the genes were up-regulated in the MD-IFX^+^ (Figure 2B). In contrast, genes associated with cell cycle pathways were down-regulated in MD-IFX^+^. From the latter observation, we reasoned that the excessive downregulated gene expression concerned cells that had undergone apoptosis. If this rationale holds true, then it is aligned with our hypothesis that mitochondrial dysfunction disrupts the proper function of macrophages to initiate inflammation-resolving processes such as clearance of apoptotic cells.

The enrichment of metabolic pathways involved the activation of the xenobiotic response system in the MD-IFX^+^ group. This includes nuclear receptors that regulate genes involved in the metabolism of xenobiotics, such as the farnesoid X receptor (FXR/*Nr1h4*) that exhibited a 1.67-fold change between the MD-IFX^+^ and WT-IFX^+^ groups causing the over-expression of the gene *Fabp6*. Moreover, the peroxisome proliferator-activated receptor *α* (*Ppara*), which regulates the expression of various UGT metabolic enzymes, ABC and SLC transporters, was also found differentially expressed. We also observed up-regulation of the peroxisome proliferator-activated receptor-gamma coactivator (PGC-1/*Ppargc1a*) gene, which interacts with PPARA and together regulate the expression of enzymes involved in the oxidation of mitochondrial fatty acids.

### MMset signal linking anti-TNF response and mitochondrial dysfunction successfully transferred to a human cohort (COLset)

We next analyzed available colon samples from a human UC cohort. Out of 21,780 quantified genes, 3,645 were differentially expressed between responders and non-responders patients. PCA showed a separation between the two groups across PC1, accompanied by an underlying intra-group variability predominantly influenced by PC2 (Figure 3A, “All DEGs”).

**Figure 3:**
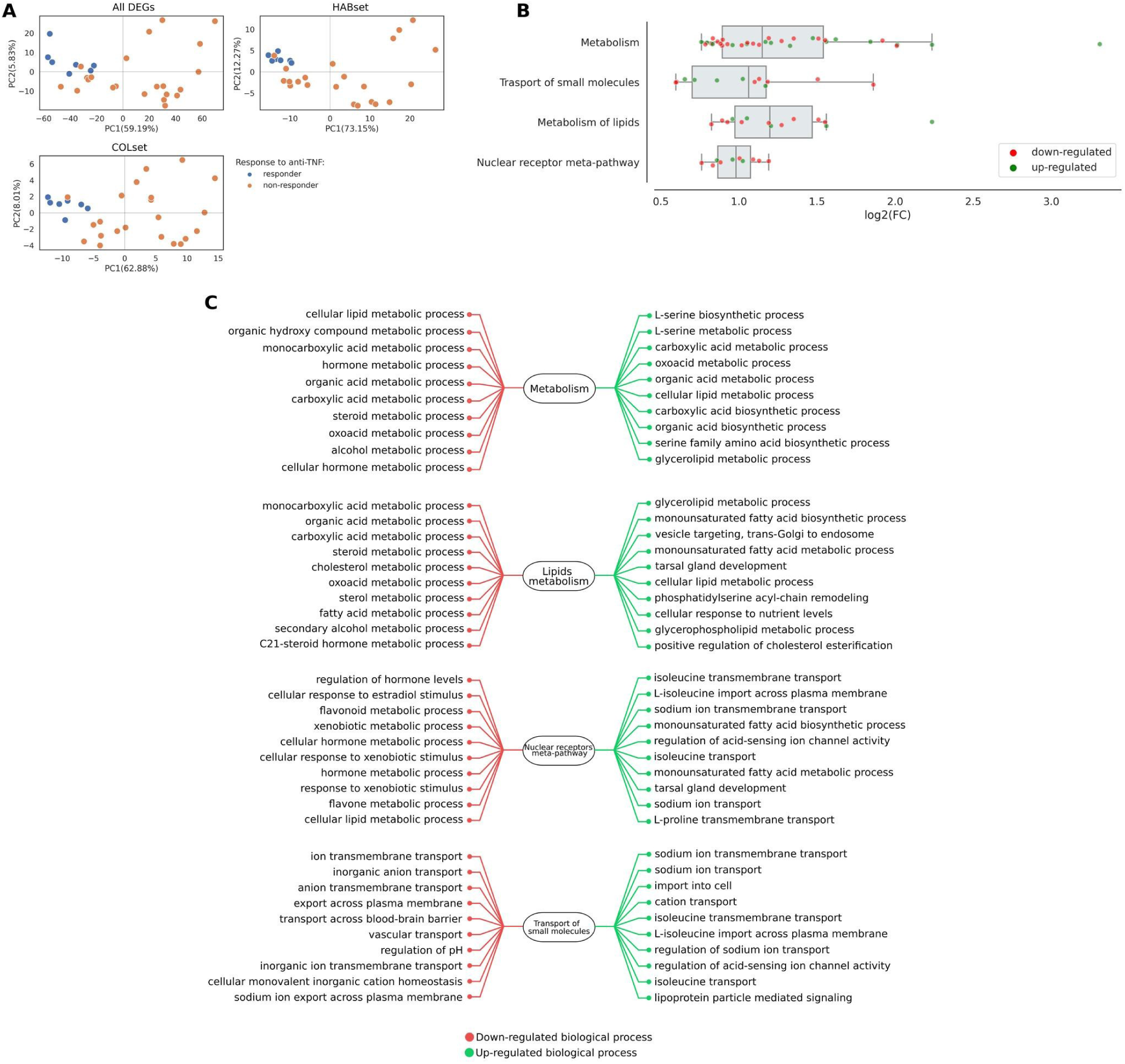
**A**) PCA using either all 3,645 genes that were differentially expressed between responders and non-responders to anti-TNF UC patients, HABset or COLset. **B)** Distribution of log2-fold changes (x-axis) between responders and non-responders to anti-TNF for each enriched pathway based on COLset. In each pathway, up and down-regulated genes are indicated for non-responders in relation to responders to anti-TNF UC patients. **C)** Up and down-regulated biological process related to each enriched pathway. Regulation is indicated for non-responders in relation to responders to anti-TNF UC patients.

We then evaluated the expression of genes that could be linked to mitochondrial dysfunction. We used the GeneCards portal [30] to query for genes that are associated with complex I. The search returned 165 genes, of which 25 were differentially expressed (supplementary materials 3). A detailed inspection revealed mitochondrial alterations, with downregulation of complex I activity components such as *MT-ND3*, *MT-ND5*, *MT-ND4*, *MT-ND4L*, *MT-TL1*, and *MT-ND2*, and up-regulation of genes such as *NLRP3*, that gets activated by excessive mitophagy [31], or *DYSF* that regulates cytosolic influx of Ca^2+^ and affects normal mitochondrial function under certain conditions [32].

To evaluate the mitochondrial aberrations that were linked to administration of anti-TNF in the murine model, we intersected the 3,645 DEGs with the MMset. This generated a subset of 127 DEGs, named COLset (supplementary materials 1). PCA showed that COLset was able to maintain the separation between responders and non-responders and increased the captured observed variance to 70% with the first two PCs (Figure 3A, “COLset”). Interestingly, although *DNAJC15* displayed statistically significant differences between responders and non-responders (p-adj=0.03), its fold-change of 1.3 between the patient groups did not allow it to be considered as differentially expressed and was not included in the COLset. This particular case exemplifies perfectly the potential pitfalls of the typical strategy followed for the discovery of DEGs, as genes that contribute to the underlying mechanism can be overlooked by strict predefined thresholds.

### COLset performs similarly to inflammation-driven gene signatures, and shows metabolism as the main enriched pathway

We then juxtaposed the discriminatory capability on the COL cohort of the COLset with a set of 89 genes from the corticosteroid-based gene signature provided recently by Haberman *et al.* [5] that we refer to as the HABset (see methods, supplementary materials 1).

Both gene sets captured above 70% of the observed variance with the first two PCs (Figure 3A), with PC1 scores showing a strong negative Pearson correlation (COLset: r=-0.69, p-value=2.6E-05, HABset: r=-0.70, p-value=2.1E-05) with *DNAJC15* mRNA levels (Figure S3). These results suggest a severity gradient of non-response that correlates with the extent of the underlying MCJ-dependent mitochondrial dysfunction. Nevertheless, both sets shared only 9 genes (*GUCA2B*, *TNIP3*, *SLC26A3*, *VSIG1*, *TRPM6*, *HMGCS2*, *MMP3*, *MMP10* and *CHP2*) suggesting that the two signatures capture different underlying gene expression regulatory components, albeit sharing functional properties.

Indeed, enrichment analysis revealed different captured pathways. The COLset enriched pathways associated with metabolism, transportation of molecules and regulation of nuclear receptors, whereas the HABset appeared linked to pathways related to the immune system, IL-17 signaling and peptide receptors (supplementary materials 2).

Finally, we evaluated the classification efficiency of the two gene sets by constructing two predictive models on the COL cohort (supplementary methods S5). The F1-score was 0.93 with the COLset and 0.97 with the HABset. As a result, the decision boundary separated well the patients groups in both cases (Figure 4). Overall, these two gene sets, although clearly different in regards to gene composition and pathway enrichment, yield similar classification results on samples from colonic tissue from UC patients undergoing anti-TNF treatment.

**Figure 4:**
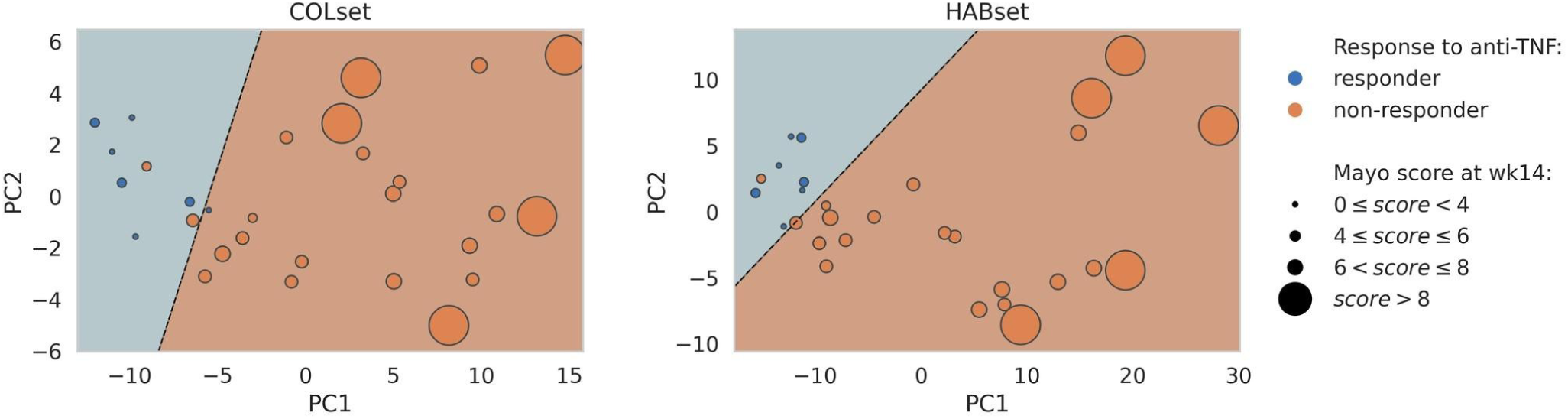
Performance of the anti-TNF logistic regression classifier based on the COL cohort and the first two PCs of the corresponding PCA that was performed using the COLset (left) or the HABset (right). Dashed line represents classifier’s decision boundary. Patients to the right side of the decision boundary (shaded orange) have been classified as non-responders by the model, whereas to the left side (shaded blue) as responders to anti-TNF. Patients have been annotated based on their observed response (color) and assigned endoscopic Mayo score (size).

### COLset classification signal was replicated in external datasets from intestinal samples of UC patients treated with anti-TNF

To validate the classification signal that was captured through COLset, we incorporated in our analysis two gene expression datasets with accession numbers GSE16879 [19] and GSE73661 [20]. In both datasets, the authors used microarrays to quantify gene expression from intestinal biopsies of UC patients before and four to six weeks after treatment with Infliximab. At baseline, we did not detect DEGs with the GSE16879 dataset, whereas 44 DEGs were identified in the GSE73661 dataset that did not show any overlap with COLset. After treatment, in each dataset we observed 805 and 1095 DEGs between responders and non-responders from which 36 and 43 DEGs were shared with COLset respectively.

We incorporated the identified DEGs in a PCA and created a classifier using the first two PCs following the same procedure as specified in supplementary methods S5. At baseline, we observed weak separation between responders and non-responders across PC2, which captured a small amount of the observed variance (Figure 5). However, after treatment, patient group separation improved drastically, and the F1-score increased when the classifier included only the DEGs that were shared with COLset. Notably, *PPARGC1A* was differentially expressed in both datasets after treatment, displaying a mean down-regulation of 1.41 FC at baseline and 2.56 FC after treatment across the two datasets regarding non-responders.

**Figure 5:**
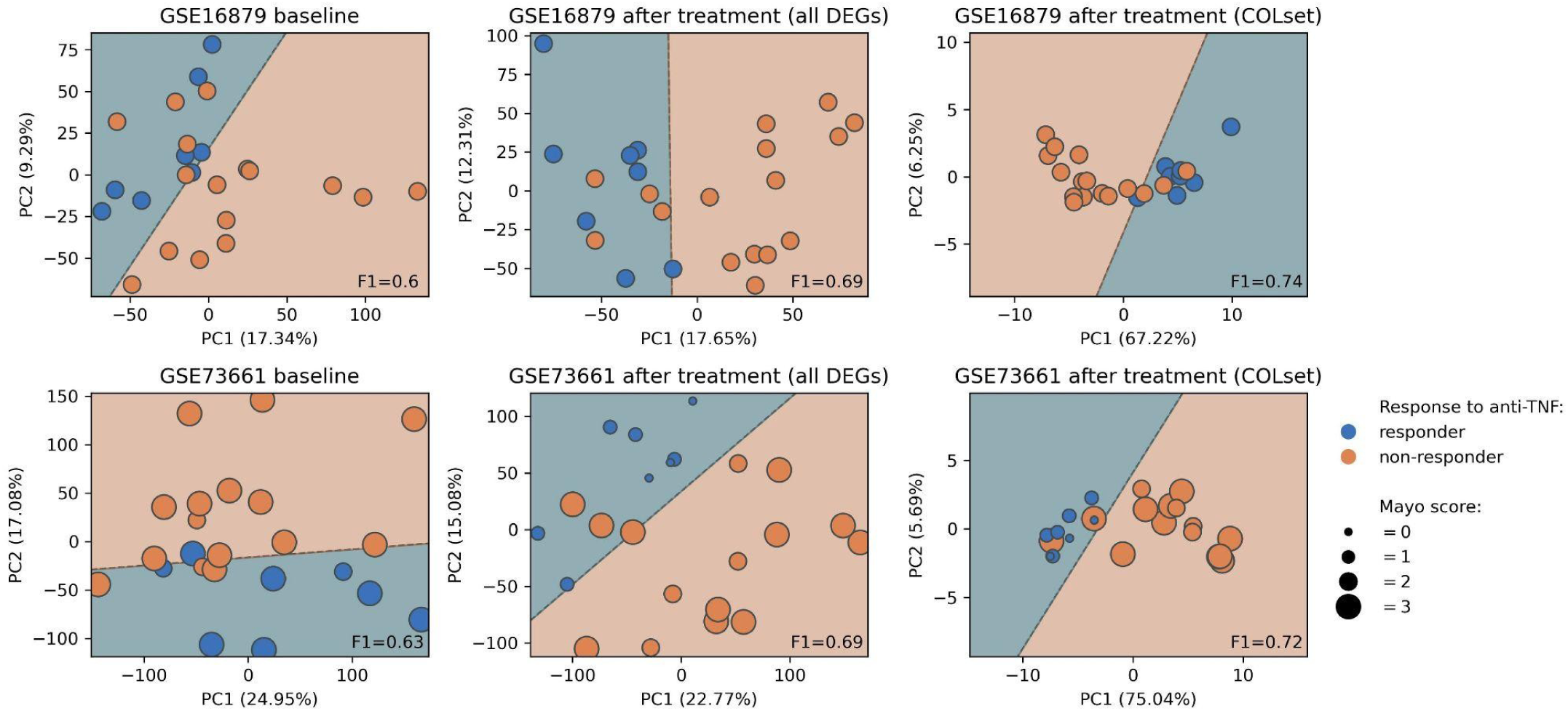
Performance of the anti-TNF logistic regression classifier based on two external dataset using the first two PCs of the corresponding PCA that was performed using gene expression at baseline, all identified DEGs after treatment and only DEGs that were common with COLset. Dashed line represents classifier’s decision boundary. Patients to the side of the decision boundary that is shaded orange have been classified as non-responders by the model, whereas to the side that is shaded blue as responders to anti-TNF. Patients have been annotated based on their observed response (color) and assigned endoscopic Mayo score (size). Mayo score was not available for the GSE16879 dataset.

Although microarrays have limited sensitivity and dynamic range compared to bulk RNASeq, these results demonstrate that the signal captured by COLset is robust across different datasets.

### COLset signal was detected in circulating blood monocytes (MONset)

We analyzed RNASeq data from peripheral blood monocyte samples with the aim to assess the persistence of the captured signal from the colon samples in the blood. We quantified 18,331 genes and identified a set of 9,262 DEGs between healthy non-IBD individuals and IBD patients with a severe disease state that were sampled right before colectomy. The intersection of the latter with the 165 genes related to complex I from GeneCards returned 83 genes (supplementary materials 3). Fifty-three genes were up-regulated (FC M=2.89, SD=1.89) and 30 were down-regulated (FC M=2.97, SD=2.15) in the IBD patients. Moreover, the IBD patients exhibited up-regulation of the majority of the NDUF genes that are associated with complex I, and down-regulation of the majority of the TMEM genes that are directly related to ATP synthesis.

To associate differential expression related to anti-TNF treatment, we intersected the 9,262 DEGs with the COLset, generating a subset of 48 genes which we named as MONset (supplementary materials 1). We attributed the narrow overlap in the homogeneous and purified nature of the samples included in the MON cohort compared to those of the COL cohort. PCA revealed a clear separation between IBD patients and non-IBD individuals, capturing 68% of the observed variance with the first two PCs (Figure 6A). Moreover, enrichment analysis showed metabolism as the sole enriched pathway encompassing 17 genes (Figure 6B-C, supplementary materials 2). These results indicate that dysregulated complex I activity can be detected in circulating monocytes, even before they migrate to colonic tissue and differentiate into macrophages.

**Figure 6:**
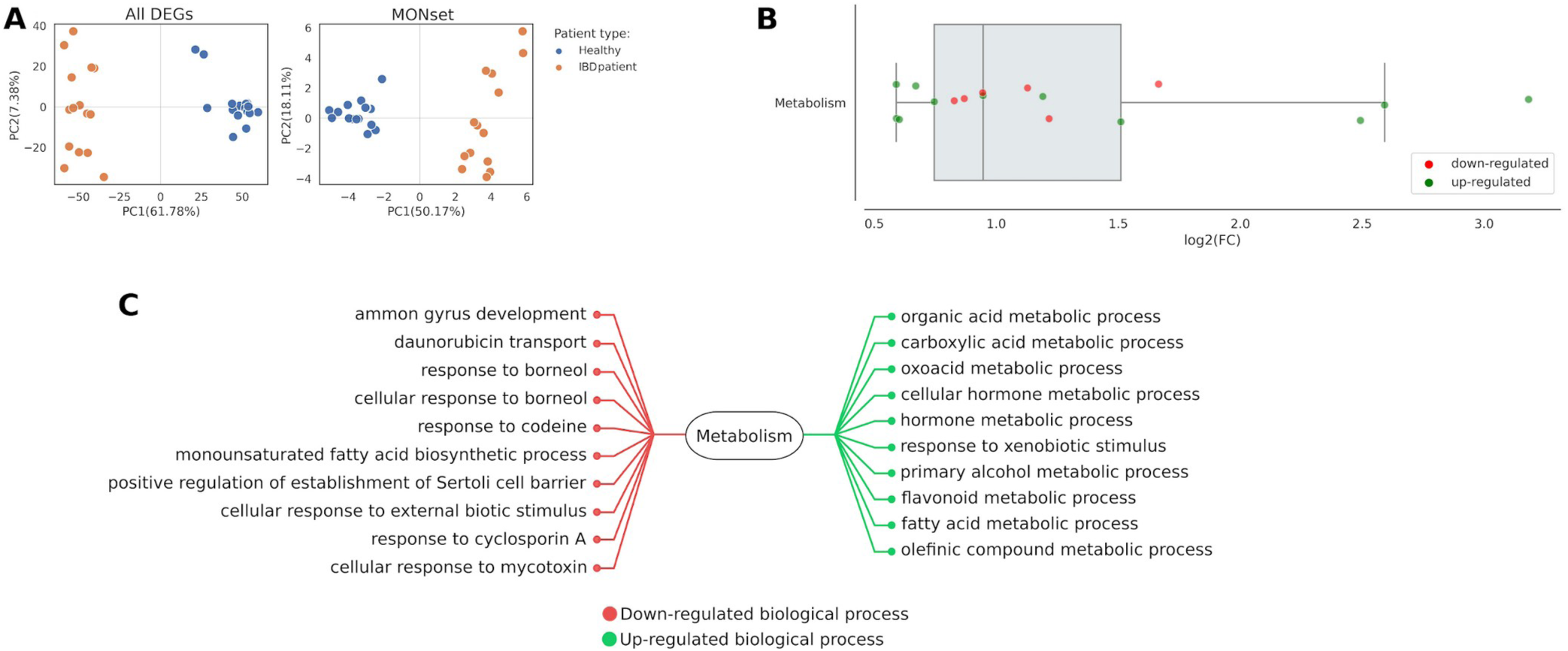
**A**) PCA using all 9262 genes that were differentially expressed between healthy and IBD patients or genes included in the MONset. **B)** Distribution of log2-fold changes (x-axis) between healthy individuals and IBD patients based on MONset. In each pathway, up and down-regulated genes are indicated for the IBD patients in relation to healthy individuals. **C)** Up and down-regulated biological process related to each enriched pathway. Regulation is indicated for IBD patients in relation to healthy individuals.

### Pre-treatment whole blood samples reveal latent classification signals for UC patient response, linked to mitochondrial dysfunction (BLDset)

Following the results from the MON cohort, our next objective was to investigate whether the classification signal of COLset, between responders and non-responders IBD patients, can also be detected in whole blood samples especially before treatment. To do so, we analyzed whole blood samples at baseline of UC patients whose status was identified at a later stage of the treatment (week-14). We quantified 24,039 genes that did not exhibit differential expression between responders and non-responders. The similarity in their transcriptomic profiles at baseline implies that unbiased classification at week-0 through classical gene signature discovery is not possible. However, non-responders displayed a mean of 1.18-fold difference (SD=0.36) compared to responders for the 165 genes associated with complex I from GeneCards (supplementary materials 3). This suggests that perturbations of mitochondrial genes are present before the initiation of the treatment.

Due to lack of differential expression in the BLD cohort, it was unfeasible to intersect the COLset with a set of DEGs in the BLD cohort. Therefore, to proceed we utilized the MONset that includes genes from the COLset that exhibit differential expression in the blood. Nevertheless, certain genes from the MONset displayed crossover trends between the BLD and MON cohorts that we attributed to the lack of responders in the MON cohort. Thus, we considered a MONset subset of 14 genes to be included in the classification process which hereafter will be referred to as BLDset (see methods section, supplementary materials 1).

During the 12-fold cross-validation of the training process, the classifier achieved a mean F1-score of 0.79 (95% CI: 0.78-0.80) with the BLD training-set, and a mean F1-score of 0.69 (95% CI: 0.59-0.79) with the BLD test-set. The panels A and B in Figure 7 depict the decision boundary applied by the classifier and the probability of the assigned class (responder or non-responder) to each included patient.

**Figure 7:**
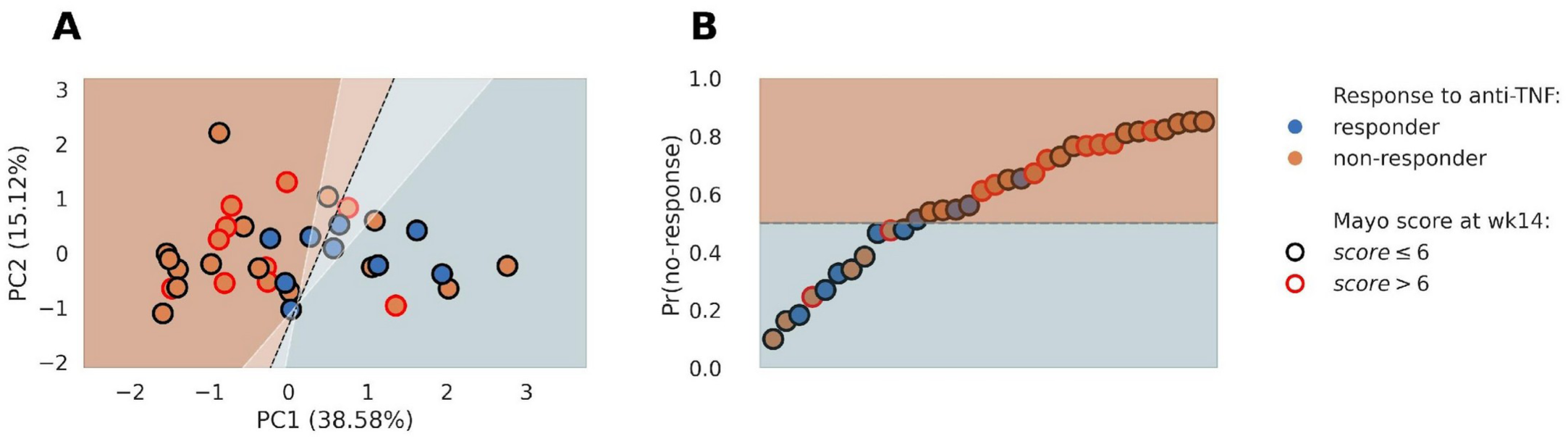
**A**) Performance of the anti-TNF logistic regression classifier based on the BLD cohort and the first two PCs of the corresponding PCA that was performed using the 14 genes of BLDset. Dashed line represents classifier’s decision boundary. White shaded area around decision boundary represents bootstrapped 95% CI, associated with the range within which the decision boundary could be positioned. Patients to the right side of the decision boundary (shaded orange) have been classified as non-responders by the classifer, whereas to the left side (shaded blue) as responders to anti-TNF. **B)** the probabilities assigned by the classifer to each patient are given, regarding the possibility of no-response to anti-TNF treatment using the BLDset. The classifier implements a probability threshold (decision boundary) of ≥50% to classify a patient as non-responder (shaded orange area) and <50% as responder to anti-TNF (shaded blue area). In all graphical representations, patients have been annotated based on observed response at week-14 after treatment (color) and assigned endoscopic Mayo score (outline).

### External replication reveals attenuated baseline BLDset classification signal in blood, with marked post-treatment improvement

To evaluate the replication of the classification process with BLDset, we utilized publicly available blood-based RNASeq data provided recently by Mishra *et al.* [24] (see methods). The study profiled two cohorts that received anti-TNF treatment (IFX and REP) and one cohort that was treated with the *α* 4*β*7 integrin inhibitor Vedolizumab (VED). Patients were followed longitudinally from baseline until week-14, and the distribution of the patient groups is given in the supplementary methods (S6).

At baseline we observed great variability of the classification performance that marked the lowest mean F1-score with the IFX cohort (Figure 8 A-B). However, the classification drastically improved after the initiation of the treatment marking the best performance on week-14. Moreover, we observed that the classification performance for the VED cohort did not follow the same pattern as with the IFX and REP cohorts after baseline, implying specificity of the BLDset towards the anti-TNF treatment.

**Figure 8:**
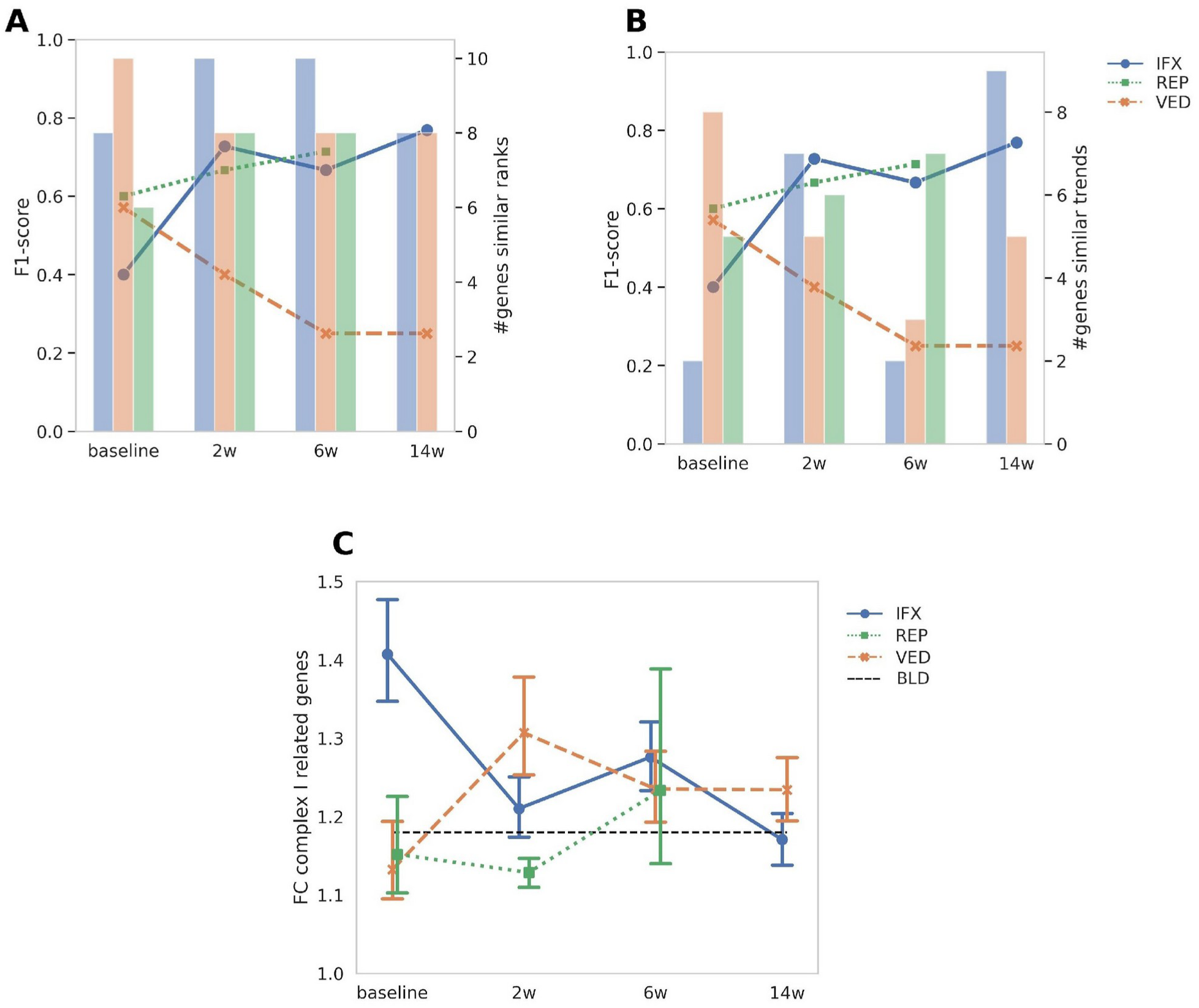
**A**) Lines represent F1-score achieved by the classifier for the cohorts that received anti-TNF (IFX, REP) or Vedolizumab (VED). Bars represent the numbers of BLDset genes from the corresponding cohorts that received the same rank as those in the BLD cohort. **B)** Lines represent F1-score achieved by the classifier for the cohorts that received anti-TNF (IFX, REP) or Vedolizumab (VED). Bars represent the number of BLDset genes from the corresponding cohorts that exhibit similar trend between patients groups when compared with those in the BLD cohort. **C)** For each cohort and time-point, the mean fold-change between the responders and non-responders UC patients is shown (error bars denote standard deviation), concerning 165 genes related to complex I. Black horizontal line represents the corresponding mean fold-changes that were observed in the BLD cohort. In all graphical representations, samples at week-14 were not available for the REP cohort.

To investigate further the variability in classification performance observed at baseline, we calculated the number of genes with matched ranks (Figure 8A) and trends (Figure 8B) between the BLDset and the external dataset, and compared them to see how they affected the performance. For the IFX cohort, we observed that it showed the highest number of matched ranks with the BLDset, but with only three genes with matched trends. This observation underscored the poor classification performance as compared to the REP and VED cohorts.

Although the above observations allowed us to explain the poor performance on the IFX cohort at baseline, the improvement in classification on this cohort at week-6 remained puzzling since the number of matched ranks and trends were relatively the same as in baseline. Initially, we observed that in both timepoints the genes *SLC18A1* and *PIF1* displayed similar trends, and the trend differences concerned the gene *EDN1* (baseline) and *STON1* (week-6) (supplementary Figure S4). We hypothesized that this observation could be associated with latent patterns of gene networks that are associated with the BLD cohort and have been indirectly incorporated in the BLDset and potentially affect the classification in the external dataset. To do so, we initially calculated a mean expression for the BLD cohort concerning the 165 complex I related genes from GeneCards. We then performed the same exercise for each cohort in the external dataset and compared them with the BLD cohort (Figure 8C). Interestingly, we observed that the performance was influenced by the captured state of the patient groups in the BLD cohort regarding these 165 complex I genes. More specifically, the performance improved in cases where the average fold-changes related to these genes between responders and non-responders were closer to those observed in the BLD cohort. Since this could be observed across all timepoints, it corroborated our hypothesis about latent gene networks that can shape the performance of a classifier.

## Discussion

Mitochondrial genes have recently been highlighted as potentially critical to explain the underlying mechanism that compromises the response to anti-TNF treatments in UC patients. Since such signals could be overshadowed by inflammation-driven pathways, we hypothesized that exploring a mouse model with a clear perturbation source is ideal to unravel obscured biology. In the current study, we first analyzed a murine model of experimental colitis characterized by mitochondrial dysfunction and lack of response to anti-TNF. To simulate a mitochondrial dysfunction, we utilized MCJ deficiency as a perturbation source which is a mitochondrial transmembrane protein that interacts with complex I [12–14]. The initial focus was to select a set of genes (MMset) that showed strong relation with mitochondrial dysregulation and response to anti-TNF treatment. Our main goal was to discover underlying expression signals associated with mitochondrial dysfunction that can be hidden by inflammatory pathways.

Notably, we also observed deficiency in MCJ expression in the COL cohort, with significant differences between the patient groups. This underscores the relevance of the mouse model with regards to the mitochondrial-driven ability to respond to anti-TNF, and we successfully transferred the classification signal captured by MMset to the COL cohort. The transferred signal (COLset) displayed a significant enrichment of metabolic pathways, and showed classification capacity similar to the recent gene signature provided by Haberman *et al.* [5] (HABset), even though the latter is enriched for inflammation pathways and both sets shared a limited number of genes. This suggests that mitochondrial genes have equal contribution to treatment outcomes as those associated with inflammation. Indeed, compromised mitochondria release damage-associated molecular patterns (DAMPs) and induce innate immune responses [31], establishing an interrelationship between both that could explain the similar classification capacity of COLset and HABset.

The replication of the classification signal of COLset in two external datasets [19,20] further corroborated the involvement of the mitochondrial regulation in the treatment response. Notably, non-responders displayed down-regulation of *PPARGC1A* both before and after treatment, with post-treatment levels becoming even more pronounced, leading to significant differential expression. This gene is a key regulator of mitochondrial biogenesis [5], and has been associated with more severe colitis and decreased complex IV activity in knock-out mice [33].

Given the restricted expression of MCJ to colonic tissue macrophages, it was pertinent to investigate whether the COLset signal extends from the colon to the bloodstream. The MON cohort allowed us to explore this aspect and we identified a significant classification signal (MONset) that was primarily enriched in metabolic pathways. Monocytes are circulating short-lived macrophage precursors that are recruited on demand from the blood to sites of inflammation such as the colon, and our results indicate that circulating monocytes in IBD patients are characterized by compromised complex I activity.

Even though classification of pre-treatment response using whole blood samples was not the main focus of this study, the results obtained by the MON cohort prompted us to explore the presence of a classification signal at baseline. Since significant insights from such an exercise can lead to valuable non-invasive classification solutions regarding response, we initially implemented a classification process using the BLD cohort and attempted to replicate the process using an external dataset. Although we did not observe robust performance at baseline, BLDset displayed specificity towards anti-TNF treatment and the whole exercise revealed the impact of gene ranks and trends on the performance across different datasets, as well as the influence of non-DEGs that are involved in the regulatory network. This kind of classification nuances could underlie some of the incongruent results seen in drug response signatures [4,6].

The BLDset was the classification signal that persisted from the colon samples to the bloodstream. Its composition, apart from *PPARGC1A*, included *PHGDH* that has been found to regulate mitochondrial homeostasis [26], and *RASGRF2* that coordinates processes related to the activity of the N-methyl-D-aspartate receptor, which has been reported to induce mitochondrial dysregulation [34]. Other members of the gene set were *SLC18A1*, a transporter with xenobiotic transmembrane activity, the detoxification gene *SULT1A1* [35], *PBLD* that is involved in the inhibition of NF-kB signaling [36] and *LRP8* and *PADI2*, which display calcium-ion activity and could be associated with neoplastic progression in UC [37].

We identified certain limitations associated with our study. The small sample size and the imbalanced patient groups limited our ability to optimize the classification process further. This limitation represents an integral part of many studies due to the difficulties of constructing a UC based cohort attributed to the low annual incidence. As a result, the incorporation of patients with Crohn’s disease (CD) represents a common workaround in order to increase statistical power. However, CD and UC patients exhibit different transcriptomic profiles [38] and following such strategies increases the risk of introducing additional latent factors into the predictive model. Furthermore, the interaction between gut microbiota and gene expression has not been incorporated into the model, since it would require integration of additional omic data such as metabolomics. There is an interplay between mitochondria and intestinal microbiota, with the latter releasing metabolites that trigger mitochondrial ROS which in turn induce an immune response [15,39]. This interaction may affect the transcriptomic profile of the patients and hence indirectly impact the performance of the classifier.

Overall, our study demonstrates the relationship between mitochondrial dysfunction and treatment response in ulcerative colitis. We showed that key regulatory networks can be masked by major inflammatory pathways during enrichment analysis, which causes critical genes to be overlooked by the stringent thresholds applied during differential gene expression. Furthermore, the patients groups displayed similar expression profiles at baseline in both colon and blood samples, complicating the identification of meaningful differences using current approaches on DEGs. We emphasize the necessity for more longitudinal studies and improved methodological approaches to better capture the expression dynamics of mitochondrial genes involved in treatment resistance and their interactions with other pathways.

## Conclusion

We highlighted the importance of mitochondrial homeostasis in relation to treatment response in the context of UC. Our results indicate that regulatory networks associated with mitochondrial dysfunction can provide promising classification signals that can be detected even in the bloodstream, leading to effective non-invasive classification solutions.

## Contributors

LA, UMM conceived and supervised the study. APC, AMC, IS, JC, JP, LGI, IRL, JOZ, MF, IMR, MG, AMA, AS, HR performed experiments and/or provided human samples. LA, JA supervised experiments. DK analyzed data. DK, UMM wrote the manuscript. All authors have reviewed the manuscript.

## Data sharing statement

The raw transcriptomic data for the mouse model and the COL, MON and BLD human cohorts are publicly available in the SRA repository under the accession number PRJNA1075662.

## Abbreviations

MM: colon samples from murine disease model
WT: all wild-type mice included in the MM dataset
WT-IFX^-^: wild-type mice without anti-TNF treatment
WT-IFX^+^: wild-type mice with anti-TNF treatment
MD: all mice of MM with MCJ-deficiency
MD-IFX^-^: mice with MCJ-deficiency and without anti-TNF treatment
MD-IFX^+^: mice with MCJ-deficiency and with anti-TNF treatment
MMset: 554 genes from MM that were differentially expressed between MD-IFX^+^ and WT-IFX^+^.
COL: colon samples at week-14 from a human cohort of responders and non-responders to anti-TNF UC patients.
COLset: 127 genes that intersected with MMset and were differentially expressed in the COL cohort.
HABset: 89 genes from Haberman *et al.* corticosteroid based gene signature that were differentially expressed in the COL cohort.
MON: monocyte samples of peripheral blood from a human cohort of healthy individuals and IBD patients.
MONset: 48 genes that intersected with COLset and were differentially expressed in the MON cohort.
BLD: whole blood samples at baseline (week-0) from a human cohort of responders and non-responders to anti-TNF UC patients.
BLDset: set of 14 genes that were used to classify response to anti-TNF from a whole blood sample at baseline.
PCA: principal component analysis
PCs: principal components
DEG: differentially expressed gene
DSS: dextran sodium sulfate

## Acknowledgements

We acknowledge all the personnel of the Hospital Clinic IBD Unit (Clinic and laboratory) who participated in patient selection and sample collection, handling and processing and we also thank all the patients who participated in this study and the Research Committee from OSI Barrualde-Galdakao.

## Supplementary Materials

**Materials 1**

List of genes that were included in MMset, COLset, HABset, MONset and BLDset.

**Materials 2**

Pathway enrichments for genes included in the MMset, COLset, HABset and MONset.

**Materials 3**

List of complex I related genes that were differentially expressed in COL and MON cohorts.

