## Supplementary Figures for "Mitochondrial dysfunction: unraveling the elusive biology behind anti-TNF response during ulcerative colitis"

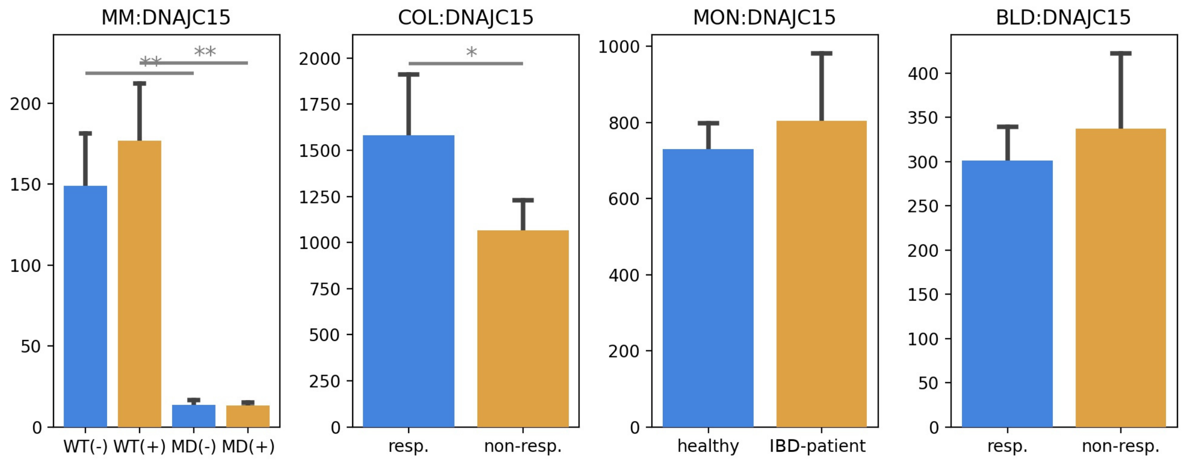


Figure S1: *Dnajc15* mRNA levels across the groups in the mouse model (MM) showing the efficiency of the MCJ deficiency process. For comparison purposes, the mRNA levels of *DNAJC15* are also shown for the COL, MON and BLD cohorts. Asterisks represent statistical significance threshold: $0.01 <$ p-adj $\leq 0.05$ (*) or p-adj $\leq 0.01$ (**). FDR-adjusted p-values (p-adj) were derived from the differential gene expression analysis with DESeq2.

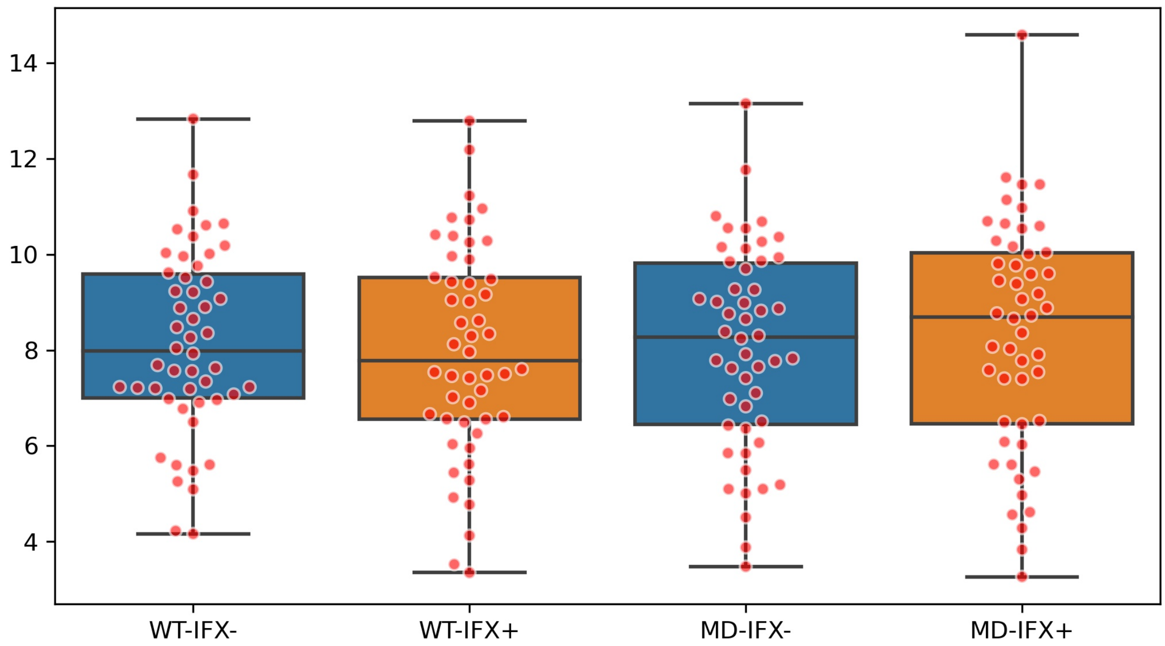


Figure S2: Boxplots of log2-transformed expression levels of 50 genes that are located on the mitochondrion for each mouse group.

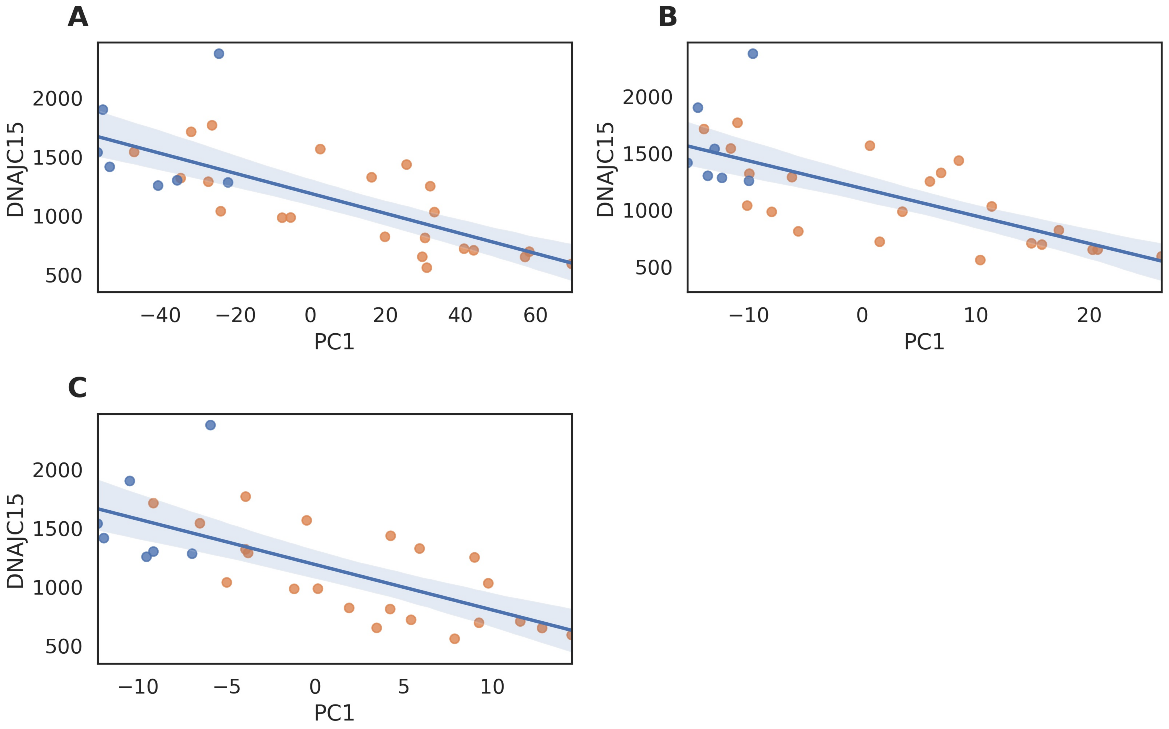


Figure S3: **A)** Pearson correlation (r=-0.73, p-value=6.80E-06) between PC1 scores of UC patients from the COL cohort, from PCA using 3,645 DEGs, and their corresponding *DNAJC15* mRNA levels (y-axis). **B)** Pearson correlation (r=-0.70, p-value=2.11E-05) between PC1 scores of UC patients from the COL cohort, from PCA using the HABset, and their corresponding *DNAJC15* mRNA levels (y-axis). **C)** Pearson correlation (r=-0.69, p-value=2.63E-05) between PC1 scores of UC patients from the COL cohort, from PCA using COLset, and their corresponding *DNAJC15* mRNA levels (y-axis).

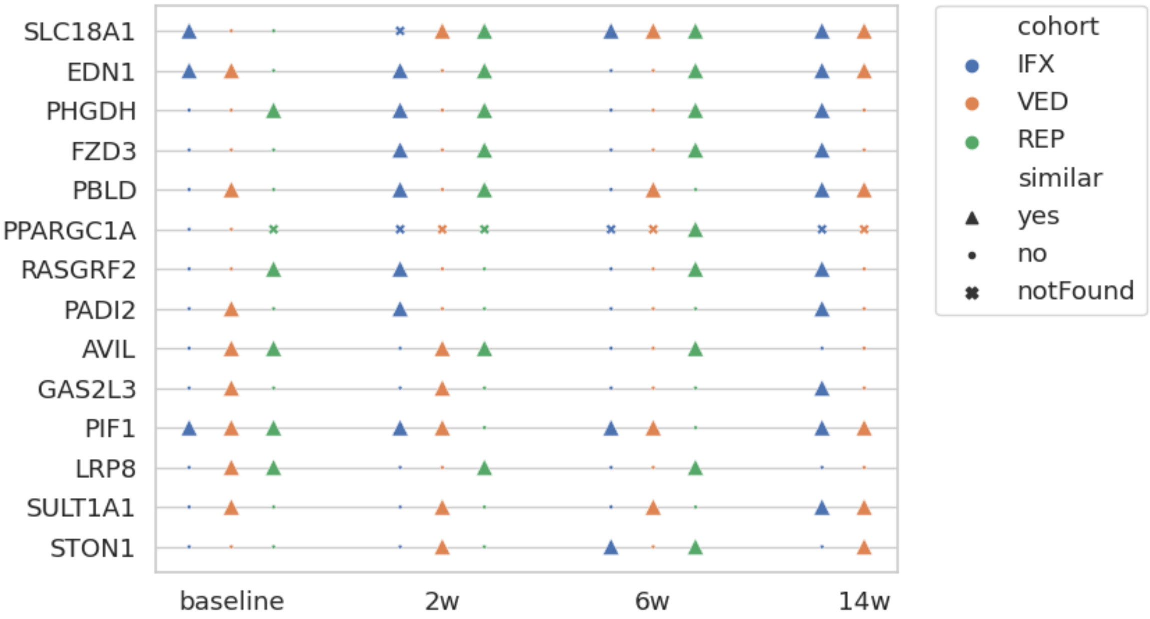


Figure S4: Genes from the BLDset with similar trends between the BLD cohort and the external dataset. Genes that were not detected in the external dataset due to their low gene expression are denoted as "notFound".
