## Supplementary Methods for "Mitochondrial dysfunction: unraveling the elusive biology behind anti-TNF response during ulcerative colitis"

### S1 Animal model and experimental design (MM)

All procedures involving animals, including their housing and care, were carried according to the guidelines of the European Union Council (Directive 2010/63/EU) and Spanish Government regulations (RD 53/2013), and with the approval of the ethics committee of CIC bioGUNE (Spain; permit number CBBA-0615) and the Competent Authority (Diputación de Bizkaia). The Animal Facility at CIC bioGUNE is accredited by AAALAC Intl.

In this study, MCJ-deficient and WT male mice on a C57BL/6 background (8-10 wk) [1] were used. Mice were maintained under specific pathogen free conditions applying standard housing conditions, in a temperature and humidity-controlled room (21-23°C temperature and 12-h light/12-h dark cycles) and were fed a standard mouse chow diet *ad libitum* (Global diet 2914, Harlam, Madison, USA).

To evaluate the response to anti-TNF treatment, colitis was promoted in wild type (WT) and MCJ deficient mice by administering dextran sodium sulfate (DSS) (36-50 kDa; TdB Consultancy) in the drinking water (2%) for 6 days followed by three-days of recovery period. Mice were divided into two groups per genotype. One group received an intraperitoneal administration of Inflectra (7.5 mg/kg Inflectra, Pfizer), an Infliximab (IFX) biosimilar proven to be safe and effective for induction and maintenance of remission in moderate to severe UC. Wild-type or MCJ deficient mice belonging to this group were denoted as WT-IFX$^{+}$ or MD-IFX$^{+}$ respectively. The second group were given saline solution where wild-type and MCJ deficient mice received the annotations WT-IFX$^{-}$ and MD-IFX$^{-}$ respectively. The mouse/human chimeric monoclonal IgG1 antibody against TNF was provided from day 3 to 6 of DSS administration. IFX was reported to also bind murine TNF [2]. The clinical progression of colitis was daily evaluated using disease activity index (DAI) score in all mice by a blind technician. DAI is the combined score of body weight loss percentage, stool consistency and the presence of gross blood in faeces. The criteria proposed by Camuesco *et al.* [3] was used to assign scores.

This murine model is referred to as MM in the manuscript. The initial distribution of mice per condition was 7 wild-type (3 WT-IFX$^{-}$ and 4 WT-IFX$^{+}$) and 8 MCJ deficient mice (4 MD-IFX$^{-}$ and 4 MD-IFX$^{+}$). In order to reduce the inter-individual variability between the mice of the same condition, we performed hierarchical clustering using normalized gene expression counts, and excluded replicates with deviating transcriptomic profile (Figure SM1). This exercise resulted in 5 wild-type (2 WT-IFX$^{-}$ and 3 WT-IFX$^{+}$) and 5 MCJ deficient mice (2 MD-IFX$^{-}$ and 3 MD-IFX$^{+}$).

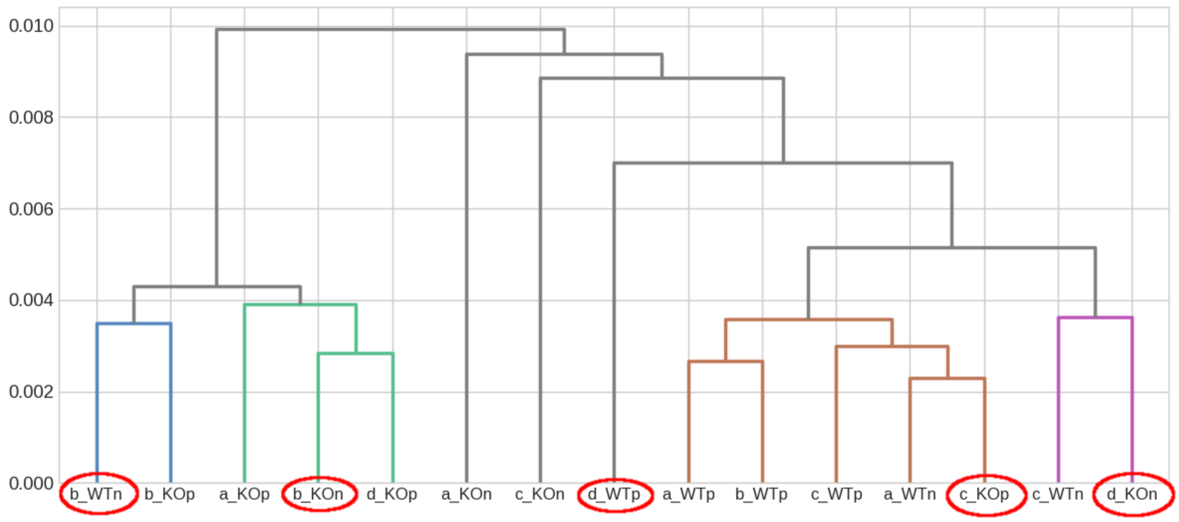


Figure SM1: Hierarchical clustering using normalized gene expression counts of the replicates WT-IFX$^-$ (WTn), WT-IFX$^+$ (WTp), MD-IFX$^-$ (KOn) and MD-IFX$^+$ (KOp). In red circle are denoted the replicates that presented deviating transcriptomic profile from the rest of the same condion and have been excluded from the downstream analysis.

### S2 Specifications for human cohorts (COL, MON and BLD cohorts)

Human cohorts are referred to with the abbreviations COL, MON and BLD. Patients in the COL and BLD cohorts were treated in the Department of Gastroenterology (Hospital CLinic de Barcelona, Spain). We obtained written consent from all patients prior to human sample processing. The samples were processed within the context of the IBD Sample Collection registered at the HCB-IDIBAPS Biobank and the protocol was approved by the Institutional Ethics Committee of the Hospital Clinic de Barcelona (HCB/2016/0389). Characteristics of the patients included in the COL and BLD cohorts are given in Table SM1.

Regarding the COL cohort, colonic biopsies were collected at week-14 after anti-TNF treatment from UC patients. Biopsies were taken at routine colonoscopies, placed in RNAlater RNA Stabilization Reagent [Qiagen] and stored at 80°C until RNA isolation. Patients were identified as responders or non-responders based on endoscopic activity.

For the MON cohort, buffy coats from human healthy donors and whole blood (15 ml) from IBD female and male patients just before colectomies were obtained from the Basque Biobank, after approval by the Clinical Research Ethics Committee from Euskadi (CEIC-E) following the Helsinki convention (PI2016034). Patient's blood samples were mixed up to 35 ml with a solution composed of phosphate-buffered saline (PBS, pH 7.2) and 2 mM EDTA to prevent coagulation. To obtain human CD14+ monocytes, blood cell suspensions were placed onto a Ficoll layer and centrifuged at 400 xg for 30 min without brake. The layer corresponding to peripheral blood monocyte cells was obtained and monocytes were positively selected using a human CD14 positive cells purification kit (Miltenyi Biotec) following the manufacturers instructions. Cells were resuspended in TRIzol (Invitrogen) and conserved at -80°C until RNA isolation.

Regarding the BLD cohort, whole blood samples were collected from UC patients into PAXgene tubes and frozen at 20°C (PreAnalytiX; Qiagen) at baseline. Patients were identified as non-responders after week-14 of anti-TNF treatment based on endoscopic activity.

**Ethnicity**

| | Caucasian | Latin American |
|---|---|---|
| non-responders | 30 | 1 |
| responders | 8 | 0 |

**week-14 MAYO score distribution**

| | 0.0 | 2.0 | 3.0 | 4.0 | 5.0 | 6.0 | 7.0 | 8.0 | 9.0 | 10.0 | 11.0 | N/A |
|---|---|---|---|---|---|---|---|---|---|---|---|---|
| non-responders | 0 | 0 | 2 | 2 | 1 | 4 | 6 | 4 | 6 | 2 | 3 | 1 |
| responders | 4 | 2 | 1 | 0 | 1 | 0 | 0 | 0 | 0 | 0 | 0 | 0 |

**Smoking status**

| | Ex-smoker | Nonsmoker | Smoker |
|---|---|---|---|
| non-responders | 10 | 18 | 3 |
| responders | 2 | 6 | 0 |

**Years from first diagnosis**

| | 0 to 10 years | 11 to 20 years | 21 to 40 years |
|---|---|---|---|
| non-responders | 9 | 9 | 13 |
| responders | 1 | 2 | 5 |

**Naive to biologics**

| | No | Yes |
|---|---|---|
| non-responders | 7 | 24 |
| responders | 1 | 7 |

**Treatment received**

| | non-responders | responders |
|---|---|---|
| Adalimumab | 4 | 0 |
| Adalimumab, Aza | 1 | 0 |
| Adalimumab, Mesalazina | 0 | 1 |
| Aza, Golimumab, Pentasa | 1 | 0 |
| Aza, Golimumab, Salofalk | 0 | 1 |
| Aza, Infliximab | 9 | 3 |
| Aza, Infliximab, Mesalazina | 3 | 0 |
| Aza, Infliximab, Mesalazina enema | 1 | 0 |
| Aza, Infliximab, Proctosteroid | 1 | 0 |
| Corticosteroids, Golimumab | 0 | 1 |
| Corticosteroids, Infliximab | 2 | 0 |
| Golimumab | 2 | 1 |
| Golimumab, Prednisona | 1 | 0 |
| Golimumab, Rifinah | 1 | 0 |
| Infliximab | 4 | 1 |
| Infliximab, Mesalazina | 1 | 0 |

Table SM1: Characteristics of patients included in the COL and BLD cohorts.

### S3 RNASeq protol and data analysis

#### S3.1 RNA extraction, library preparation and sequencing

RNA concentration was measured using Qubit 2.0, with Qubit RNA assay kit (Invitrogen, Cat.# Q32855). RNA integrity was assessed by Agilent 2100 Bioanalyzer, using Agilent RNA 6000 Pico or Nano Chips (Agilent Technologies, Cat.# 5067-1513 or Cat.# 5067-1511, respectively). Sequencing libraries were prepared following TruSeq Stranded mRNA Sample Preparation Guide (Part # 15031058 Rev. E) using the TruSeq® Stranded mRNA Library Prep kit (Illumina Inc. Cat.# 20020594) and TruSeq RNA CD Indexes (Illumina Inc. Cat.# 20019792) (MM libraries) or TruSeq RNA Single Indexes (Illumina Inc. Cat.# 20020492 and 20020493) (MON, COL, BLD libraries).

Briefly, starting from 1000 ng (MM libraries) or 100 ng of total RNA (MON, COL, BLD libraries), mRNA was purified, fragmented and primed for cDNA synthesis. cDNA first strand was synthesized with SuperScript-II Reverse Transcriptase for 10 min at 25°C, 15 min at 42°C, 15 min at 70°C and pause at 4°C. cDNA second strand was synthesized with Illumina reagents at 16°C for 1 hour. Then, A-tailing and adaptor ligation were performed. Finally, enrichment of libraries was achieved by PCR (30 sec at 98°C; 15 cycles of 10 sec at 98°C, 30 sec at 60°C, 30 sec at 72°C; 5 min at 72°C and pause at 4°C). In the case of BLD samples, some modifications to this protocol were needed, before first strand cDNA synthesis, globin mRNA depletion was carried out using QIAseq FastSelect-globin kit (Qiagen, Cat.# 334377) and 17 cycles of PCR were performed for library enrichment.

Afterwards, libraries were visualized on an Agilent 2100 Bioanalyzer using Agilent High Sensitivity DNA kit (Agilent Technologies, Cat.# 5067-4626) and quantified using Qubit dsDNA HS DNA Kit (Thermo Fisher Scientific, Cat.# Q32854).

All libraries were sequenced using Illumina platforms to obtain a minimum of 20 million 100 bp paired-end reads (MM libraries) or 27.5 million 50 bp single-end reads (MON, COL, BLD libraries).

#### S3.2 Preprocessing of raw RNASeq data and enrichment analysis

Initially, RNASeq reads with length $< 20$ after adapters removal or trimming based on Phred33 quality $< 20$ were removed with CUTADAPT (v1.18) [4]. Subsequently, reads that mapped rRNA sequences were filtered with SortMeRNA (v2.1) [5] and the remaining reads from the mouse model were mapped to the GRCm38 mouse reference with STAR (v2.6) [6], whereas the human reference GRCh38.p13 was used for the human samples. From the aligned reads, those that were not uniquely mapped or had mapping quality $< 20$ were removed with SAMTOOLS (v1.11) [7] and the remaining reads were used for gene quantification with HTSeq (v0.11) [8].

For each cohort, genes with count $< 10$ across all samples were removed and the remaining raw gene counts were normalized to correct library size differences, serving as a basis for differential gene expression analysis with DESeq2 [9]. For visualization purposes, such as principal component analysis (PCA) or hierarchical clustering, the gene counts were regularized log-transformed with DESeq2. A gene was characterized as differentially expressed (DEG) for a given condition contrast, if it met the following criteria: FDR-corrected p-value (p-adj) $\leq 0.05$ and absolute fold-change $|FC| \geq 1.5$.

Enrichment analysis of DEGs was performed using the Cytoscape plug-in ClueGO [10] where databases from KEGG, Reactome and WikiPathways were queried for pathway enrichment and annotation. On the other hand, enriched biological processes were queried using the

ToppCluster portal [11]. From the resulted annotations, only statistically significant enriched pathways were considered (p-adj $\leq 0.05$).

#### S4 Construction of BLDset and the concept of trends

The classification capacity of a set of genes can be influenced by two factors: the trends and the ranks of the included genes. The term rank refers to the relative position of a gene in the set after arranging the mRNA levels of the included genes in descending order. The term trend refers to the relationship (high or low) of the mRNA levels of a given gene in a patient group when compared to another group. Based on these definitions, change in rank indicates change on the impact of a given gene, whereas change in trend (crossover trend) implies the existence of one or more moderator variables. The latter in a simplistic manner are usually referred to as cofounding factors. However, since there is a possibility genes that were not captured by differential expression analysis to act as moderators, changes in trends should be evaluated with caution and contextual awareness.

For the construction of the BLDset we considered 14 genes from the MONset that displayed the same trend between responders and non-responders in the COL and BLD cohorts. To do so, we defined a function that accepts mRNA levels of a given gene as argument and returns 0, when the mean mRNA levels of the responders is smaller than the mean levels of the non-responders, or 1 otherwise. Finally, after considering only patients from the BLD cohort for whom samples were available in the COL cohort, the BLDset included those genes of MONset for which the function returned the same value when presented with the corresponding mRNA levels from the COL and BLD cohorts (Equation 1). Therefore, genes with crossover trend between the BLD and COL cohorts were excluded. The described filtering of the BLD patients was implemented only for the creation of the BLDset and not for the construction of the classifier where all patients from the BLD cohort were considered.

Finally, we performed the same process with HABset (Equation 1) and as a result we included an additional set of 18 genes increasing the total number of genes in BLDset to 32.

$$
\begin{aligned}
f(g) &= \begin{cases} 0, & \bar{g}_{\text{responders}} < \bar{g}_{\text{non-responders}} \\ 1, & \bar{g}_{\text{responders}} > \bar{g}_{\text{non-responders}} \end{cases} \\
\text{BLDset} &= \{g \in \text{MONset} \mid f(COL_{(g)}) = f(BLD_{(g)})\}
\end{aligned} \tag{1}
$$

We incorporated BLDset into a classifier that utilized the BLD cohort as a training-set after regressing out age and sex and stabilized gene expression variance with DESeq2. To avoid overfitting and multicollinearity, we performed PCA on the BLD cohort and based a non-regularized logistic regression classifier (Equation 2) with Scikit-learn [12] on the first two PCs and used the weights 0.3 and 0.7 for the non-responder and responder classes respectively so as to mitigate the impact of the imbalanced conditions.

$$
\begin{aligned}
Pr(\text{response} = 1|\text{BLDset}) &= logistic(\beta_0 + \beta_1 PC1_{BLD} + \beta_2 PC2_{BLD} + \epsilon) \\
\text{response} &= \begin{cases} 1, & \text{no-response to anti-TNF} \\ 0, & \text{response to anti-TNF} \end{cases}
\end{aligned} \tag{2}
$$

### S5 Construction of a classifier for response to anti-TNF at week-14 using the COL cohort.

The aim of this classifier was to evaluate the discriminatory capacity of the COLset against the corticosteroid based gene signature provided by Haberman *et al.* [13] which is referred to as HABset in the manuscript.

From the COL cohort, we regressed out age and sex and stabilized gene expression variance with DESeq2. To avoid overfitting and multicollinearity, we performed PCA and based the classifier on the first two principal components (PCs). Using the COLset and HABset sequentially, we carried out a non-regularized logistic regression (Equation 3) with Scikit-learn [12] where we used the weights 0.3 and 0.7 for the non-responder and responder classes respectively so as to mitigate the impact of the imbalanced conditions. Finally, the performance of the two gene sets was assessed using the F1-score.

$$\begin{gathered} Pr(\text{response} = 1|\text{COLset}) = logistic(\beta_0 + \beta_1 PC1_{COL} + \beta_2 PC2_{COL} + \epsilon) \\ Pr(\text{response} = 1|\text{HABset}) = logistic(\beta_0 + \beta_1 PC1_{COL} + \beta_2 PC2_{COL} + \epsilon) \\ \text{response} = \begin{cases} 1, & \text{no-response to anti-TNF} \\ 0, & \text{response to anti-TNF} \end{cases} \end{gathered} \tag{3}$$

### S6 Patient groups distribution in the external dataset

In Table SM2 the distribution of the patient groups is given for the UC cohorts provided by Mishra *et al.* [14] (GSE191328) and used for replication of the classification process. Two cohorts received anti-TNF treatment (IFX and REP) and one cohort was treated with the $\alpha 4\beta 7$ integrin inhibitor Vedolizumab (VED).

| Cohort | Timepoint | N (non-resp./resp.) |
|---|---|---|
| IFX | baseline | 9 (6/3) |
| | week-2 | 8 (5/3) |
| | week-6 | 9 (6/3) |
| | week-14 | 9 (6/3) |
| REP | baseline | 9 (6/3) |
| | week-2 | 9 (6/3) |
| | week-6 | 9 (6/3) |
| VED | baseline | 10 (4/6) |
| | week-2 | 9 (3/6) |
| | week-6 | 8 (3/5) |
| | week-14 | 8 (2/6) |

Table SM2: Distribution of patient groups of the UC cohorts provided by Mishra *et al.* [14] (GSE191328).

[11] Vivek Kaimal, Eric E Bardes, Scott C Tabar, Anil G Jegga, and Bruce J Aronow. Toppcluster: a multiple gene list feature analyzer for comparative enrichment clustering and network-based dissection of biological systems. *Nucleic acids research*, 38(suppl_2): W96–W102, 2010.

[12] F. Pedregosa, G. Varoquaux, A. Gramfort, V. Michel, B. Thirion, O. Grisel, M. Blondel, P. Prettenhofer, R. Weiss, V. Dubourg, J. Vanderplas, A. Passos, D. Cournapeau,

M. Brucher, M. Perrot, and E. Duchesnay. Scikit-learn: Machine learning in Python. *Journal of Machine Learning Research*, 12:2825–2830, 2011.

[13] Yael Haberman, Rebekah Karns, Phillip J Dexheimer, Melanie Schirmer, Judith Somekh, Ingrid Jurickova, Tzipi Braun, Elizabeth Novak, Laura Bauman, Margaret H Collins, *et al.* Ulcerative colitis mucosal transcriptomes reveal mitochondriopathy and personalized mechanisms underlying disease severity and treatment response. *Nature communications*, 10(1):1–13, 2019.

[14] Neha Mishra, Konrad Aden, Johanna I Blase, Nathan Baran, Dora Bordoni, Florian Tran, Claudio Conrad, Diana Avalos, Charlot Jaeckel, Michael Scherer, *et al.* Longitudinal multi-omics analysis identifies early blood-based predictors of anti-tnf therapy response in inflammatory bowel disease. *Genome medicine*, 14(1):1–20, 2022.
